# Discovery of the TGCB microprotein translated from the lncRNA CHASERR

**DOI:** 10.64898/2025.12.04.692433

**Authors:** Bernardo Bonilauri, Xiaochun Yang

**Affiliations:** Stanford Cardiovascular Institute, Stanford University School of Medicine, Stanford, CA 94305, USA; Department of Medicine, Stanford University School of Medicine, Stanford, CA 94305, USA

**Author notes:** Correspondence to: Bernardo Bonilauri, Stanford Cardiovascular Institute, Department of Medicine, Stanford University, 240 Pasteur Drive, Stanford, CA 94304, USA.

**Keywords:** microprotein, lncRNA, translation, noncoding, small open reading frame

## Abstract

Long noncoding RNAs (lncRNAs) are key regulators of gene expression, and an increasing number have been found to contain small open reading frames (smORFs) that can be translated into functional microproteins. *CHASERR* is a conserved cis-regulatory lncRNA positioned upstream of *CHD2*, where it fine-tunes *CHD2* dosage. Deletions spanning the *CHASERR* gene were recently identified in children with a syndromic, early-onset neurodevelopmental disorder, underscoring the clinical importance of this regulatory locus. Here, we report that *CHASERR* is not strictly noncoding but encodes a previously unannotated 41-amino-acid microprotein, which we name TGCB. Integrative analyses of bulk and single-cell transcriptomics, 47 public Ribo-seq datasets, coding-potential predictors, and multispecies alignments support active translation and evolutionary constraint of the TGCB smORF. Public mass-spectrometry data provide reproducible peptide-spectrum matches corresponding to its N-terminal region, and immunofluorescence confirms nuclear and cytoplasmic localization. Structural prediction reveals a compact α-helical microdomain flanked by intrinsically disordered regions, characteristic of regulatory microproteins. Notably, all pathogenic deletions affecting the *CHASERR–CHD2* region remove the entire TGCB coding sequence, raising the possibility that loss of this microprotein contributes to the associated neurodevelopmental phenotypes. These findings position *CHASERR* as a bifunctional transcript and uncover a conserved microprotein embedded within a clinically relevant regulatory locus.

## Introduction

*CHASERR* is a relatively conserved cis-regulatory long noncoding RNA (lncRNA) positioned upstream of the *CHD2* gene, where it acts as a critical dosage regulator. *CHASERR* and *CHD2* show tightly coordinated expression across tissues in mouse and human, and mechanistic studies have demonstrated that *CHASERR* functions strictly in cis to restrain *CHD2* output[1]. Disruption of this regulation leads to a pronounced increase in Chd2 mRNA and protein levels, with severe biological consequences: homozygous *Chaserr* loss causes early postnatal lethality, and heterozygous deficiency results in marked growth impairment in mice, underscoring the developmental sensitivity of this regulatory pair[1].

The clinical relevance of this axis has been highlighted by recent reports of three unrelated children presenting with early-onset neurodevelopmental delay and encephalopathy caused by heterozygous de novo deletions encompassing *CHASERR*[2]. Although the deletions differed in size (22 kb, 8.4 kb, and 25 kb), all removed at least the promoter and first three exons of *CHASERR*. Induced pluripotent stem cells (iPSCs) derived from two of these patients exhibited significantly elevated CHD2 protein levels compared with sex-matched controls, providing compelling human evidence—consistent with mouse models— that *CHASERR* haploinsufficiency leads to *CHD2* overexpression[2].

In this *Short Communication*, we leverage integrative analyses of bulk and single-cell transcriptomics, ribosome profiling (Ribo-seq), comparative genomics, structural prediction, cellular localization, and public proteomics to demonstrate that *CHASERR* is not strictly noncoding but instead encodes a conserved 41–amino–acid microprotein, TGCB, translated from a smORF within exon 1. This discovery places *CHASERR* within the expanding class of bifunctional transcripts. Importantly, the existence of TGCB introduces a previously unrecognized molecular component at the *CHASERR*–*CHD2* locus—one that may contribute to the dosage sensitivity and neurodevelopmental phenotypes observed in individuals carrying deletions of this region. By revealing this additional layer of regulatory complexity, our findings expand the conceptual framework and open new avenues for mechanistic and clinical investigation into *CHD2*-related neurodevelopmental disorders.

## Results and Discussion

To obtain a global view of *CHASERR* expression across human tissues, we analyzed GTEx v10 and quantified its median expression levels. *CHASERR* was most highly expressed in tissues of the central and peripheral nervous systems, followed by the ovary, lung, thyroid, and arterial structures (**Figure 1A**). A *CHASERR*–gene correlation analysis across all GTEx tissues identified *CHD2* as one of the strongest correlates (ρ_s_ = 0.80), fully consistent with the known regulatory relationship between the two loci (**Figure 1B**).

**Figure 1.**
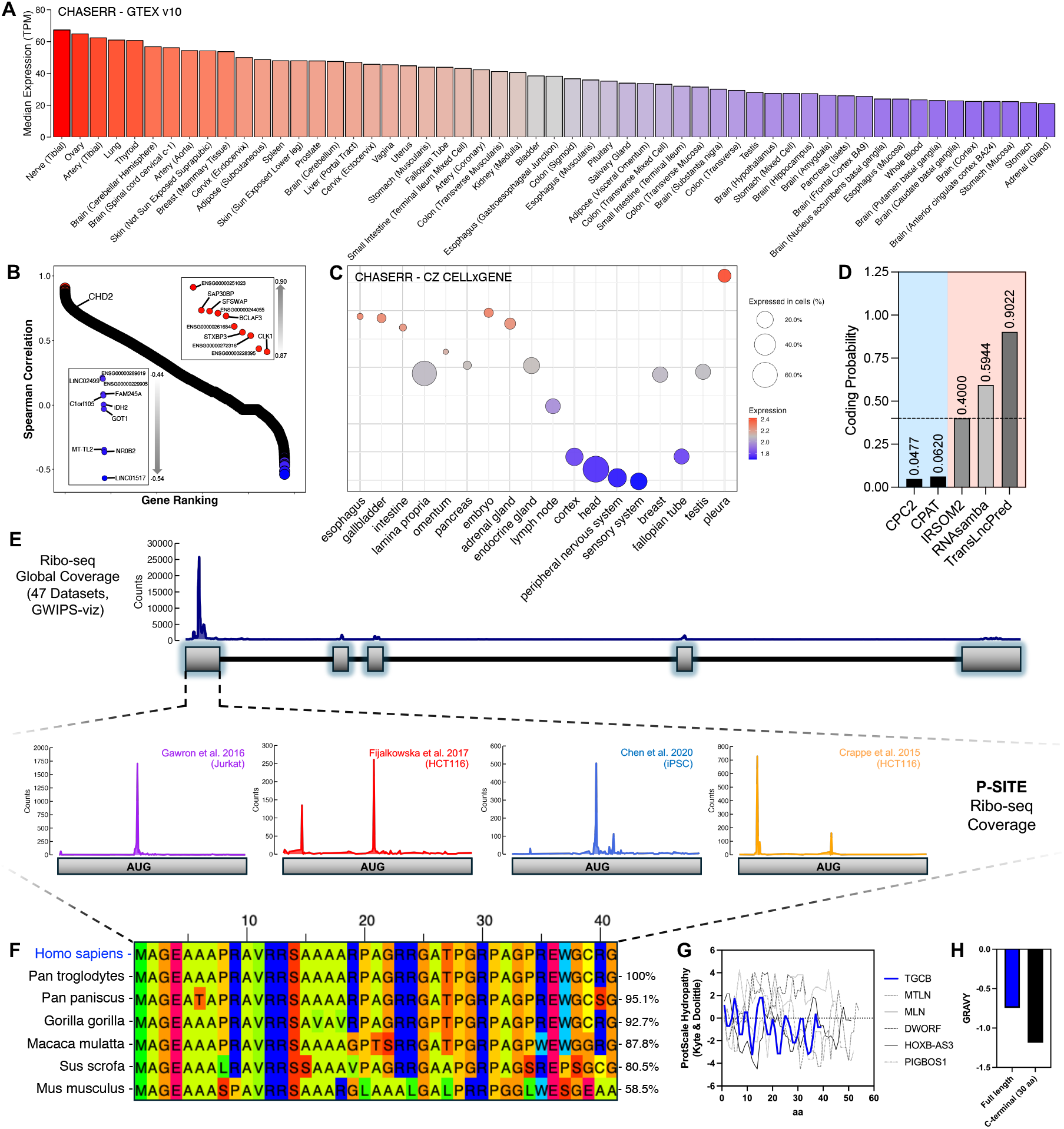
Genomic context, expression landscape, and translational evidence for *CHASERR* and its encoded microprotein TGCB. **(A)** Median expression of *CHASERR* across human tissues (GTEx v10). **(B)** *CHASERR* co-expression analysis based on Spearman rank correlation across human transcriptomes. Zoomed view of the top positively and negatively correlated genes are shown. **(C)** Single-cell RNA-seq expression of *CHASERR* across major human cell types and tissues using the CZ CELLxGENE meta-atlas. Dot size reflects the percentage of expressing cells; color indicates normalized expression levels. **(D)** Coding-potential predictions for the *CHASERR* gene using five independent software (CPC2, CPAT, IRSOM2, RNAsamba, TransLncPred). **(E)** Global Ribosome profiling (Ribo-seq) coverage from 47 publicly available datasets (GWIPS-viz), revealing accumulation of ribosomal footprint in the exon 1 of *CHASERR*. Lower panels shows Ribo-seq P-site peak precisely aligned to the start codon (AUG) of the smORF; representative individual datasets confirming reproducible ribosomal occupancy in diverse cell types (i.e., Jurkat, HCT116, and iPSCs). **(F)** Multispecies alignment of the TGCB microprotein across mammals demonstrating high evolutionary conservation. **(G)** Hydropathy analysis of the TGCB microprotein (blue line) compared with several documented microproteins (gray lines). TGCB exhibits an overall hydrophilic profile, including a marked drop in hydropathy across its C-terminal region. **(H)** GRAVY index of the full-length TGCB microprotein (blue) and of its C-terminal 30 amino acids (black), highlighting the strongly hydrophilic character of both the global sequence and its terminal domain.

Because *CHASERR* expression is widespread, we next examined single-cell RNA-seq datasets from CZ CELLxGENE[3]. *CHASERR*-positive cells were abundant throughout the brain, head, peripheral nerves, and sensory systems. Additional *CHASERR*-expressing populations were detected in the pleura and other tissues (**Figure 1C**), particularly within immune-related lineages, including lymphocytes, phagocytes, myeloid leukocytes, monocytes, macrophages, dendritic cells, and neutrophils.

Several lncRNAs have recently been shown to contain ribosome-associated small open reading frames (smORFs) that undergo active translation, producing microproteins with defined cellular functions and revealing an additional, previously overlooked layer of lncRNA biology[4–8]. To assess whether *CHASERR* might share this feature, we evaluated its coding potential using five independent computational tools; three of them supported the presence of a putative coding region within *CHASERR* (**Figure 1D**). In addition, IRSOM2[9]—designed to identify bifunctional RNAs by rejecting strictly coding and strictly noncoding classifications—predicted *CHASERR* to exhibit a bifunctional signature.

Motivated by these results, we analyzed ribosomal footprints from 47 public Ribo-seq datasets available on GWIPS-viz[10,11]. The aggregated data revealed clear ribosomal occupancy in exon 1, including a sharp initiating P-site peak precisely at the predicted AUG start codon—strong evidence of active translation (**Figure 1E**). Consistent with our observations, a recent preprint also noted Ribo-seq signal over *CHASERR* exon 1 in both human and mouse datasets, even though its primary aim was to investigate *CHASERR* regulation by targeting motifs in the last exon that induce formation of a *CHASERR–CHD2* fusion transcript. Notably, this fusion incorporates the complete *CHD2* coding sequence and can be translated into full-length CHD2 protein[12].

Comparative phylogenetic analysis showed that the predicted *CHASERR*-encoded microprotein is highly conserved across mammals, with strong amino-acid identity extending throughout higher primates (**Figure 1F**). In humans and the primate species analyzed, the smORF spans 126 nucleotides (41 amino acids), whereas orthologous ORFs in pig and mouse are substantially longer—306 nucleotides (101 aa) and 354 nucleotides (117 aa), respectively. Based on these results, we designate this 41-amino-acid microprotein as TGCB (i.e., Translated Gene *CHASERR* Bicistronic).

Although pervasive translation of noncanonical smORFs often generates unstable peptides with hydrophobic C-terminal tails destined for BAG6-mediated triage and proteasome degradation[5,13,14], TGCB displays the opposite biochemical signature. Strikingly, both the Kyte–Doolittle hydropathy profile and the GRAVY index revealed a strongly hydrophilic composition (GRAVY < 0), and the final ~30 C-terminal residues exhibited strongly negative hydropathy values (**Figure 1G and 1H**). These features are consistent with a translated microprotein with biochemical stability and potential functional relevance.

To predict the subcellular localization of TGCB, we first applied DeepLoc 2.1[15], which indicated a soluble microprotein with high probability of nuclear localization, with a secondary likelihood of cytoplasmic localization (**Figure 2A**). We then expressed TGCB in cells and examined its distribution by immunofluorescence. Consistent with the computational prediction, TGCB was detected predominantly in the nucleus and cytoplasm (**Figure 2B**). To further corroborate the existence of TGCB at the protein level, we mined publicly available mass spectrometry datasets through OpenProt 2.0[16], which curates proteomics evidence for noncanonical ORFs. We queried the human alternative protein entry annotated within *CHASERR* (41 aa) and retrieved the corresponding annotated spectra for manual inspection. Notably, across three independent datasets, we identified peptide–spectrum matches mapping to the N-terminal TGCB peptide (AGEAAAPR), with consistent b- and y-ion series supporting sequence assignment (**Figure 2C**). The PSM confidence values reported by OpenProt (PSM scores of 0.36, 0.29 and 0.10) are modest in absolute terms, which is expected for short tryptic peptides derived from microproteins. Nevertheless, the recurrence of the same peptide across independent studies, together with reproducible fragment-ion support, provides orthogonal evidence consistent with the translation of the predicted TGCB microprotein.

**Figure 2.**
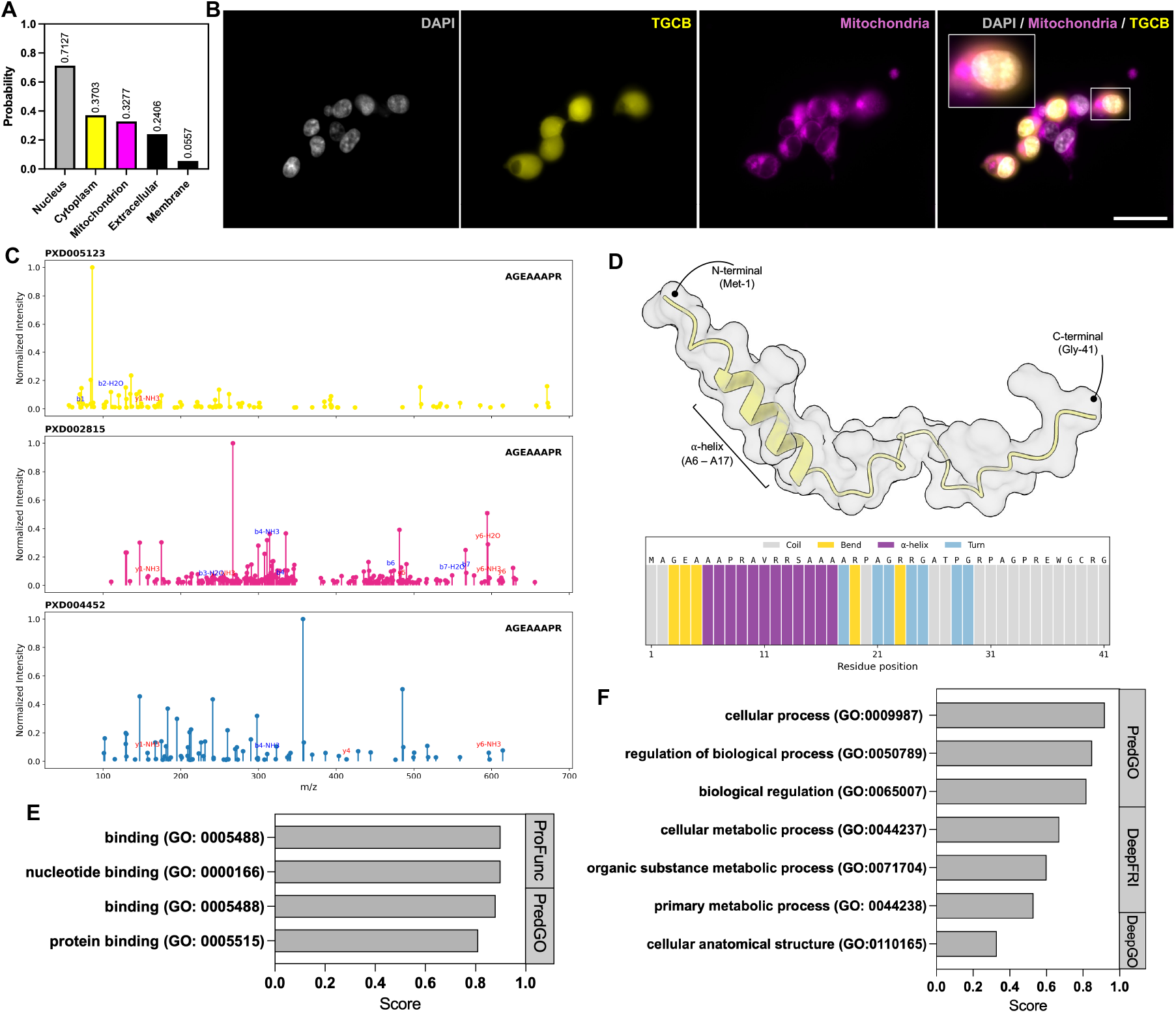
Discovery and molecular properties of the TGCB microprotein. **(A)** Subcellular localization prediction generated with DeepLoc2.1. **(B)** Immunofluorescence of cells overexpressing TGCB; nuclei are shown in grey, TGCB microprotein in yellow, and mitochondria in magenta. Scale bar = 20 µm. **(C)** Annotated MS/MS spectra from three independent public datasets (PXD005123, PXD002815, and PXD004452) showing peptide–spectrum matches (PSMs) corresponding to the N-terminal TGCB tryptic peptide AGEAAAPR. **(D)** AlphaFold2 structural model of TGCB highlighting the compact α-helical core. Linear secondary-structure representation derived from DSSP, showing a single α-helix (A6–A17) flanked by intrinsically disordered regions. **(E)** Predicted Molecular Function terms generated from consensus computational analysis. **(F)** Predicted Biological Process Gene Ontology terms for TGCB (DeepFRI, ProFunc, PredGO, DeepGO).

To gain insight into the structural organization of TGCB, we predicted its three-dimensional fold using AlphaFold2[17]. The top-ranked model revealed a compact α-helical core spanning residues A6–A17, flanked by flexible N– and C–terminal regions consistent with intrinsic disorder (**Figure 2D**). This structural arrangement was further supported by per-residue secondary-structure assignments obtained using DSSP, which identified a single α-helical segment within the central region of the protein, while classifying the remaining residues predominantly as coil or turn. Together, these results indicate that TGCB adopts a partially structured architecture composed of a short stable helix embedded within disordered tails—a common organizational motif among regulatory microproteins.

Finally, we assessed TGCB’s potential functional signatures using four orthogonal protein-function predictors (DeepFRI[18], DeepGO[19], ProFunc[20], PredGO[21]). All tools returned only broad and low-specificity Gene Ontology terms—most consistently enriched for binding, protein binding, nucleotide binding, cellular process, and biological regulation (**Figure 2E and F**). Putative protein interaction partners of TGCB remain unknown and will require unbiased proteomic mapping. This pattern, common among microproteins lacking conserved domains, is consistent with TGCB functioning as a small regulatory cofactor rather than an enzyme, in line with its structural architecture consisting of a single α-helix flanked by intrinsically disordered regions.

The findings presented here are not only clinically relevant to the neurodevelopmental disorders and epilepsy observed in children carrying *CHASERR* deletions[2] but also point to broader disease contexts in which *CHASERR* dysregulation may play a role. For instance, recent studies have reported significantly elevated *CHASERR* levels in the blood of patients with Alzheimer’s disease compared to healthy individuals[22]. Similarly, *CHASERR* is markedly upregulated in glioma tissue relative to normal tissue, and its high expression correlates with poorer overall survival in glioma patients[23]. In mouse and rat models, *Chaserr* upregulation has been observed in the brain following middle cerebral artery occlusion and reperfusion, as well as in primary cortical neurons subjected to oxygen-glucose deprivation and reoxygenation, respectively[24]. Moreover, *CHASERR* becomes highly methylated in human bronchial epithelial cells upon exposure to influenza virus, suggesting transcriptional modulation in response to infectious or inflammatory stimuli[25]. Together, these observations highlight *CHASERR* as a disease-responsive regulatory node across diverse pathological settings and underscore the importance of understanding the contribution of its newly discovered microprotein, TGCB, to these biological and clinical phenotypes.

### Limitations of the study

Although this *Short Communication* provides convergent genomic, ribosome-profiling, structural, localization, and proteomic evidence supporting translation of the *CHASERR*-encoded microprotein TGCB, several limitations remain. First, the absence of a high-affinity, sequence-validated anti-TGCB antibody currently precludes definitive confirmation of endogenous protein expression. Second, the intrinsic challenges of detecting microproteins—short length, rapid turnover, and low abundance—limit the sensitivity of conventional proteomic approaches. Finally, the functional relationship between TGCB, *CHASERR*, and *CHD2* is not yet defined, and mechanistic studies will be required to determine whether TGCB contributes directly to *CHD2* dosage control or to broader processes. Despite these constraints, the evidence presented here provides a strong foundation and highlights the need for focused follow-up studies using endogenous tagging strategies, CRISPR-based perturbations, and high-sensitivity proteomics. Our aim is to bring TGCB to the attention of the scientific community and catalyze deeper investigation into its regulatory and clinical significance.

## Materials and Methods

### GTEX and CELLxGENE expression analysis

Median tissue-level expression values for *CHASERR* were obtained from GTEx[26] v10 (RNA-SeQC version 2.4.2). The median TPM matrix was imported into R and filtered by gene annotation to extract *CHASERR* expression across tissues. Expression values were reshaped into long format and ranked to generate a genome-wide expression profile. To assess co-expression patterns, Spearman correlation coefficients were computed between *CHASERR* and all other GTEx-annotated genes across tissues. Correlation values were used to identify the highest positively and negatively associated genes.

Single-cell RNA expression patterns were analyzed using aggregated transcriptomic profiles from CZ CELLxGENE (meta-atlas release)[3]. The dataset provided tissue-cell-level *CHASERR* expression and the proportion of cells expressing the gene within each tissue. *CHASERR*-expressing tissues and cellular populations were quantified and visualized using dot plots encoding expression magnitude and cell-type representation.

### Coding probability prediction

Coding potential was assessed using five complementary prediction tools, each based on distinct computational principles. CPC2[27] employs a prediction model derived from four intrinsic sequence features to discriminate coding from noncoding transcripts. CPAT[28] uses a logistic regression classifier trained on four sequence-derived attributes to estimate coding probability. IRSOM2[9] implements a self-organizing map algorithm capable of identifying bifunctional RNAs with both coding and noncoding characteristics. RNAsamba[29] applies a neural network–based classification approach to infer coding potential from sequence composition. TransLncPred[30] integrates multiple translationally relevant features using an XGBoost classifier optimized to distinguish coding regions within long noncoding transcripts. For all tools, we adopted a coding probability cutoff of ≥ 0.4.

### Ribo-seq analysis

Ribosome profiling coverage tracks were obtained from GWIPS-viz[10,11], which compiles and aggregates publicly available Ribo-seq datasets. BigWig files corresponding to global ribosomal footprints and initiating ribosome tracks were downloaded and converted to bedGraph format using the hg38 reference genome. Coverage values across the *CHASERR* locus were extracted and visualized in R using custom scripts to generate position-resolved density plots. These aggregated ribosomal footprints were then used to assess translation signatures over exon 1 and to examine P-site alignment around the predicted AUG start codon of the TGCB smORF.

### Lentiviral expression construct and production

The smORF encoding the human TGCB microprotein (41 aa) was synthesized de novo (abm®, Richmond, Canada) and sequence-verified. The insert contained a standard Kozak consensus sequence and was subcloned by the manufacturer into the pLenti-III-CMV-GFP expression vector, which carries a puromycin resistance cassette, generating a CMV-driven TGCB–GFP fusion construct. High-titer recombinant lentiviral particles were produced by abm® using a second-generation packaging system. Viral supernatants were purified, concentrated, and titrated by functional infectious-unit assays, and supplied at ≥107 IU/mL.

### Lentiviral transduction

HEK293 cells were maintained in DMEM high glucose supplemented with 10% fetal bovine serum (FBS) and 1% penicillin–streptomycin at 37 °C and 5% CO_2_. For transduction, cells were seeded the day prior to infection at approximately 50% confluence in 24-well plates. Viral supernatant was added directly to the cells at the desired multiplicity of infection (MOI 1–5) in the presence of 8 µg/mL polybrene (Santa Cruz Biotechnology, #sc-134220). Plates were centrifuged at 800 × *g* for 1 h at 32 °C and returned to the incubator at 37 °C, 5% CO_2_. After 12–16 h, viral medium was replaced with fresh complete growth medium, and cells were allowed to recover for 48–72 h before downstream assays.

### Immunofluorescence microscopy

Transduced cells expressing TGCB–GFP were plated on glass-bottom dishes and imaged under live-cell conditions. Mitochondria were labeled using MitoTracker™ Deep Red FM (Invitrogen™, #M22426) according to the manufacturer’s protocol, and nuclei were counterstained with DAPI.

### Mass spectrometry data analysis

Public proteomics evidence for TGCB was queried through OpenProt 2.0[16], which reports peptide-spectrum matches (PSMs) supporting noncanonical ORFs. We retrieved all PSMs associated with the CHASERR-encoded 41-aa microprotein and downloaded the corresponding .mgf spectra. Spectra were visualized using a custom Python script that normalized peak intensities and overlaid theoretical b– and y–ion fragment series for the predicted N-terminal peptide AGEAAAPR.

### AlphaFold2 structure prediction

TGCB microprotein structure prediction was performed using AlphaFold2[17] (version 2.3.2) with default parameters and no structural templates. Five models were generated, and the top-ranked model—selected by pLDDT and PAE scores—was used for analysis. pLDDT values were uniformly high within the α-helical core (residues 6–17) and low in the N- and C-terminal tails, consistent with intrinsic disorder.

### Secondary-structure assignment

Per-residue secondary-structure classifications were extracted from the AF2-predicted PDB using DSSP (version 4.5.3). DSSP codes (H, G, I, E, B, T, S, C) were mapped to a residue-level linear visualization using a custom Python script that aligned the TGCB amino-acid sequence to its DSSP labels.

### Microprotein function prediction

To infer potential molecular functions and biological processes associated with the TGCB microprotein, we performed a consensus computational analysis using four independent protein-function prediction tools: DeepFRI[18], DeepGO[19] (version 1.0.26), ProFunc[20], and PredGO[21]. Each method applies a distinct algorithmic framework—graph convolutional networks (DeepFRI), deep-learning–based ontology classifiers (DeepGO), structural and sequence-derived feature mapping (ProFunc), and integrative GO-prediction models (PredGO)—allowing for orthogonal assessment of putative functional signatures. Results from all predictors were parsed, harmonized, and ranked in R using custom scripts that grouped Gene Ontology (GO) terms and extracted the top-scoring functional categories.

### CRediT authorship contribution statement

Bernardo Bonilauri: Writing – review & editing, Writing – original draft, Visualization, Validation, Project administration, Methodology, Investigation, Formal analysis, Conceptualization. Xiaochun Yang: Validation – review & editing.

